# HLA-DR4Pred2: an improved method for predicting HLA-DRB1*04:01 binders

**DOI:** 10.1101/2023.07.24.550447

**Authors:** Sumeet Patiyal, Anjali Dhall, Nishant Kumar, Gajendra P. S. Raghava

**Affiliations:** Department of Computational Biology, Indraprastha Institute of Information Technology, Okhla Phase 3, New Delhi-110020, India

**Keywords:** HLA-DRB1*04:01, BLAST, Immunotherapy, Machine Learning, Web server

## Abstract

HLA-DRB1*04:01 is associated with many disease that include sclerosis, arthritis, diabetes and Covid19. Thus, it is important to scan binders of HLA-DRB1*04:01 in an antigen to develop immunotherapy, vaccine and protection against these diseases. One of the major limitations of existing methods for predicting with HLA-DRB1*04:01 binders is that these methods trained on small datasets. This study present a method HLA-DR4Pred2 developed on a large dataset contain 12676 binders and equal number of non-binders. It is an improved version of HLA-DR4Pred, which was trained on a small dataset contain only 576 binders and equal number of binders. All models in this study were trained, optimized and tested on 80% of data called training datasets using five-fold cross-validation; final models were evaluated on 20% of data called validation/independent dataset. A wide range of machine learning techniques have been employed to develop prediction models and achieved maximum AUC of 0.90 and 0.87 on validation dataset using composition and binary profile features respectively. The performance of our composition based model increased from 0.90 to 0.93 when combined with BLAST search. In addition, we also developed our models on alternate or realistic dataset that contain 12676 binders and 86300 non-binders and achieved maximum AUC 0.99. Our method perform better than existing methods when we compare the performance of our best model with performance of existing methods on validation dataset. Finally, we developed standalone and online version of HLA-DR4Pred2 for predicting, designing and virtual scanning of HLA- DRB1*04:01(https://webs.iiitd.edu.in/raghava/hladr4pred2/; https://github.com/raghavagps/hladr4pred2).

**Key Points:**

- HLADR4Pred2.0 is an update of HLADR4Pred
- Predict the binding or non-binding peptides for MHC-Class II allele HLA- DRB1*04:01
- Used alignment free and alignment based hybrid approach
- Motifs which are highly specific to HLA-DRB1*04:01 binders
- Benchmark the performance of the other existing methods with HLADR4Pred2.0

**Author’s Biography:**

1. Sumeet Patiyal is currently working as Ph.D. in Computational biology from Department of Computational Biology, Indraprastha Institute of Information Technology, New Delhi, India
2. Anjali Dhall is currently working as Ph.D. in Computational Biology from Department of Computational Biology, Indraprastha Institute of Information Technology, New Delhi, India.
3. Nishant Kumar is currently working as Ph.D. in Computational biology from Department of Computational Biology, Indraprastha Institute of Information Technology, New Delhi, India
4. Gajendra P. S. Raghava is currently working as Professor and Head of Department of Computational Biology, Indraprastha Institute of Information Technology, New Delhi, India.

## Introduction

The human leukocyte antigen (HLA) complex is a highly polymorphic genomic region located at chromosome 6 in the human genome[1–3]. The HLA system is classified into three major categories I, II and III, where I (HLA-A, -B, -C) and II (HLA-DP, -DQ, -DR) genes are polymorphic in nature [4]. IMGT/HLA is the largest repository of HLA related sequences report thousands of human major histocompatibility complex associated alleles and genomic sequences[5–7]. HLA class-I molecules display intracellular peptides to CD8+ T cells whereas HLA class-II molecules composed two polypeptide chains (α and β) and presents extracellular peptides to CD4+ T cells. HLA-class II alleles mainly presented on antigen presenting cells for instance, B cells, macrophages, DCs, etc.[8–10]. The binding groove of MHC-II molecules is open from both sides which enables long length peptides to enlarge the binding grooves from the flanking regions as shown in Figure 1 [11,12]. Majority of MHC class II alleles present peptides that are derived from pathogenic proteins [13,14]. MHC class II alleles carry a peptide and present it on the cell surface, where it interacts with T cell receptors (Figure 1C). This interaction activates CD4+ T cells, which secrete cytokines such as interferon-gamma, tumor necrosis factor, and granulocyte-macrophage colony-stimulating factor (GM-CSF) to initiate immune responses.

**Figure 1:**
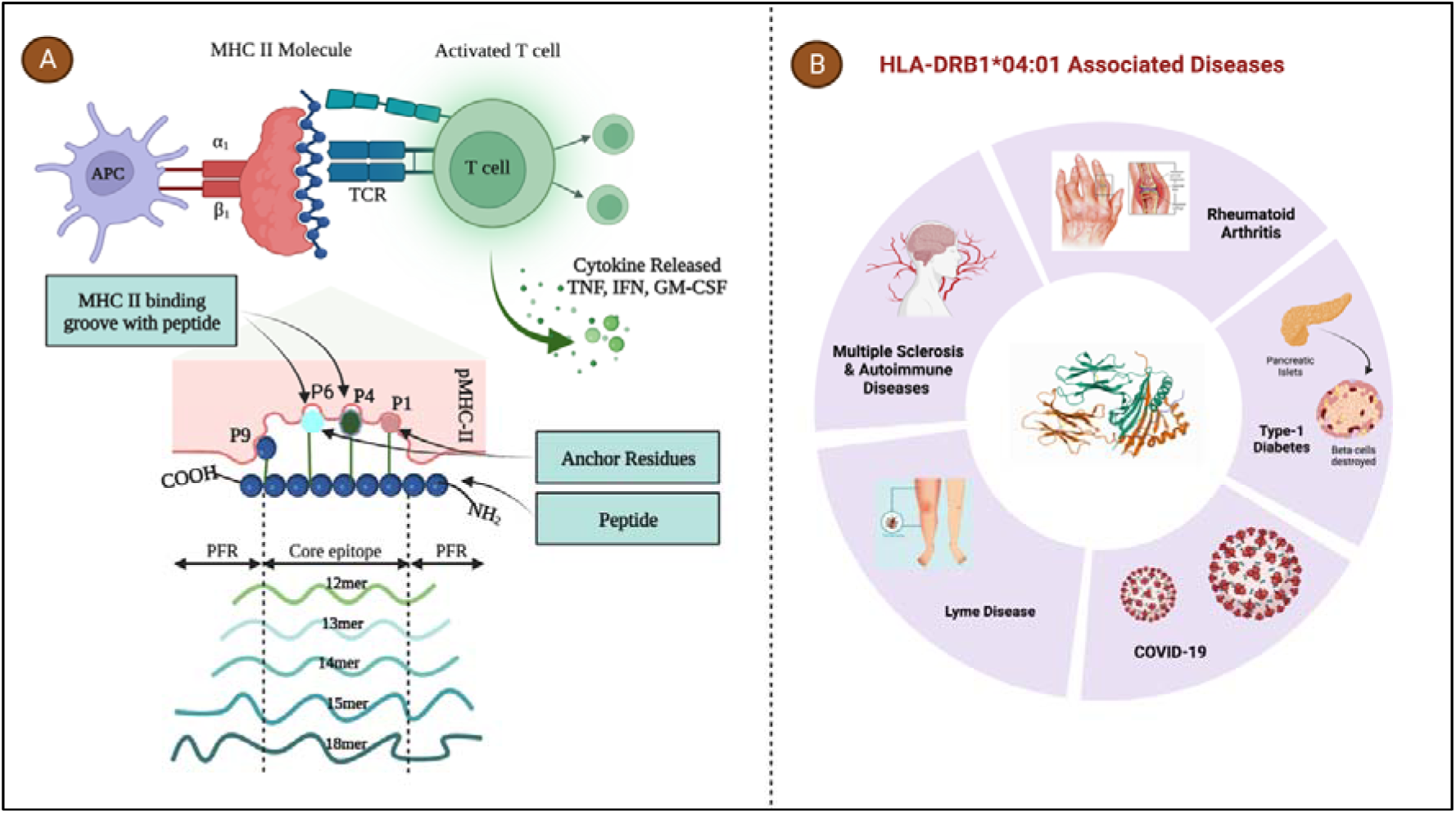
(A) Pictorial representation of peptide conformations presented by MHC-II molecules, anchor residues of peptides bound to the allele-specific pockets of MHC-II; (B) Association of HLA-DRB1*04:01 allele with number of diseases

In the past, several studies have shown that the HLA-DR4 gene is highly correlated with several diseases [15–18], especially HLA-DRB1∗04:01 is associated with the development of multiple sclerosis [19,20], autoimmune disorders (AID) [21], type 1 diabetes [22], Lyme disease [23], COVID-19 severity, and rheumatoid arthritis [24]. HLA-DR4 molecules plays a significant role in autoimmune disorders initiation and progression. Therefore, it is of utmost importance to determine the epitopes which bind to HLA-DRB1*04:01 in order to understand or cure several autoimmune disorders [25–29]. Studies also reveal that patients positive with HLA-DR4 associated alleles have maximum chances of having autoimmune disorders therefore it could be a significant as genetic biomarker. Researcher developed a number of experimental techniques for the detection of HLA-peptide bindings, but they are time-exhaustive and cost-effective [30,31]. Virtual scanning of HLA-DR4 binders at genome level is not practically feasible due to the time, cost, and complexity involved in experimental techniques. A number of computational methods have been developed to predict HLA-DR4 binders, which can facilitate the scientific community in performing scanning of binders at large scale [32–36]. In 2004, our group developed the HLA-DR4Pred method, which is widely used and cited by the scientific community. One of the major limitations of HLA- DR4Pred is that it is trained on a small dataset that was available in 2004.

In this study, we present an upgraded version of HLA-DR4Pred, which was trained using the largest dataset available in the IEDB database. We deployed state-of-the-art techniques to enhance the accuracy of predicting HLA-DRB104:01 binders. Initially, we created two datasets: the main dataset containing 12,676 HLA-DRB104:01 binders and an equal number of non-binders, and the realistic dataset containing 12,676 binders and 86,300 non-binders. In order to classify binders and non-binders, we developed classification models using a wide range of machine learning techniques. All models were evaluated using internal and external validation techniques. It is known fact that similar sequences or patterns have similar functions, this facts have been utilized in this study to improve accuracy of binder prediction. In this study, we used BLAST for sequence similarity search and MERCI software for motif/pattern discovery and searching. The culmination of our work is an ensemble method that combines machine learning and similarity-based approaches, enabling the precise prediction of HLA-DR binders. For a comprehensive understanding of our study’s workflow, please refer to Figure 2.

**Figure 2:**
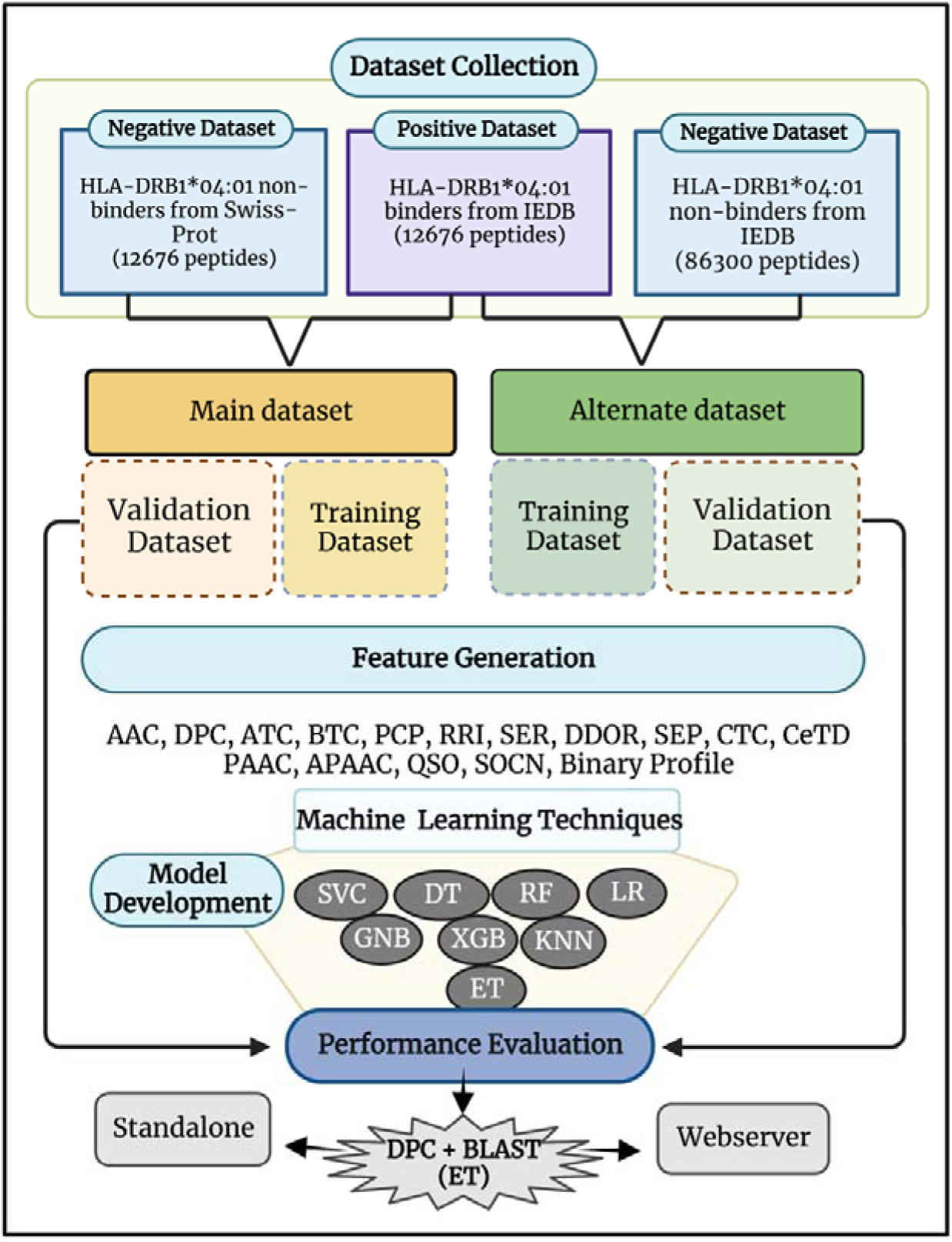
Overall workflow of algorithm implemented in this study

## Material and methods

### Dataset Creation and Preprocessing

In this study, we have extracted experimentally validated HLA-Class II allele HLA- DRB1*04:01 binding peptides from the immune epitope database (IEDB) [37]. Initially, the total number of binding peptides extracted from IEDB was 19,665, with lengths varying from 8 to 32 amino acids. After eliminating identical peptides and peptides containing non-natural amino acids, we were left with 12,880 unique peptides. Further analysis of the peptide lengths revealed that 98.4% (i.e., 12,676 peptides) had lengths ranging from 9 to 22 amino acids (as depicted in Supplementary Figure 1). Consequently, we selected these 12,676 peptides to constitute our positive dataset. In such prediction methods, one of the primary challenges is acquiring an experimentally validated negative dataset, which consist of non-binders of HLA-DRB1*04:01 allele. We generated peptides of length 9 to 22 residues from protein in Swiss-Prot database. We randomly picked up 12,676 peptides from Swiss-port generated peptides and assigned them as non-binders. Finally, we create a main dataset that contain of 12676 experimentally validated binders obtained from IEDB and 12676 non-binders generated from proteins in Swiss-Prot.

In addition to main dataset, we also generated an alternate dataset or realistic dataset that contain 12676 binders and 86300 non-binders obtained from IEDB. These 86300 non-binders were obtained from IEDB; after removing HLA-DRB1*04:01 binders and peptides do not have length between 9-22 amino acids. Both the dataset was further divided into training and validation dataset, where 80% of the data constitute training and the remaining 20% make validation dataset. To avoid the biasness in the length distribution in training and validation dataset, we have arranged all peptides as per their length and then transferred every fifth peptide into the validation dataset and rest constitutes training dataset.

### Composition Analysis

To check the abundance of each amino acid in each dataset, we have calculated the composition of each amino acid using equation 1 have utilized the amino acid composition module of Pfeature to calculate the composition of positive and negative set separately in each dataset.

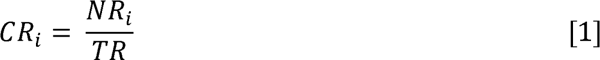

Where, CR_i_ represents composition of residue i; NR_i_ is total number of residues of type i; and TR stands for total number of residues.

### Position Conservation Analysis

In order to explore the position specific preference of residues, we have created the logos using the software named ‘Weblogo’ [38] webserver. A stack of amino acids that are measured in bits is graphically represented by this software. The overall height of each stack represents the sequence conservation at that position. On the other hand, the height of symbols within a stack reflects the relative frequency of the relevant amino or nucleic acid at that particular position [38]. Ensuring the peptide fix length parameter is a prerequisite for creating a logo. Since, the minimum length of the considered peptide is 9, hence to achieve the fix length criteria we have taken the first 9 residues from the N-terminal and last 9 residues from the C-terminal. Finally, by concatenating both the regions we have create a fix length peptide with 18 residues. We have created the “weblogo” for HLA-DRB1*04:01 binders and non-binders in the, main, and alternate dataset.

### Generation of Features

In this study to represent the sequence as a numerical vector, we have utilized the composition and binary profile module of Pfeature [39]. By using this tool, we have computed an extensive set of features, including composition and binary profile-based features. Using composition module we have calculated fifteen different type of features such as AAC, APAAC, ATC, BTC, CeTD, CTC, DDOR, DPC, PAAC, PCP, QSO, RRI, SER, SEP, and SOCN. By implementing binary profile-based module, we have computed the four different features such as binary profile of first nine residues (N_9_), binary profile of last nine residues (C_9_), and combination of N_9_ and C_9_ binary profile (N_9_C_9_). In Supplementary Table 1, we have reported the length of the vector size generated by composition, and binary profile based features. In Table 1, we have shown the example sequences of different length and highlighted the regions in the sequences which is designated as N_9_, C_9_ and N_9_C_9,_ respectively.

**Table 1:**
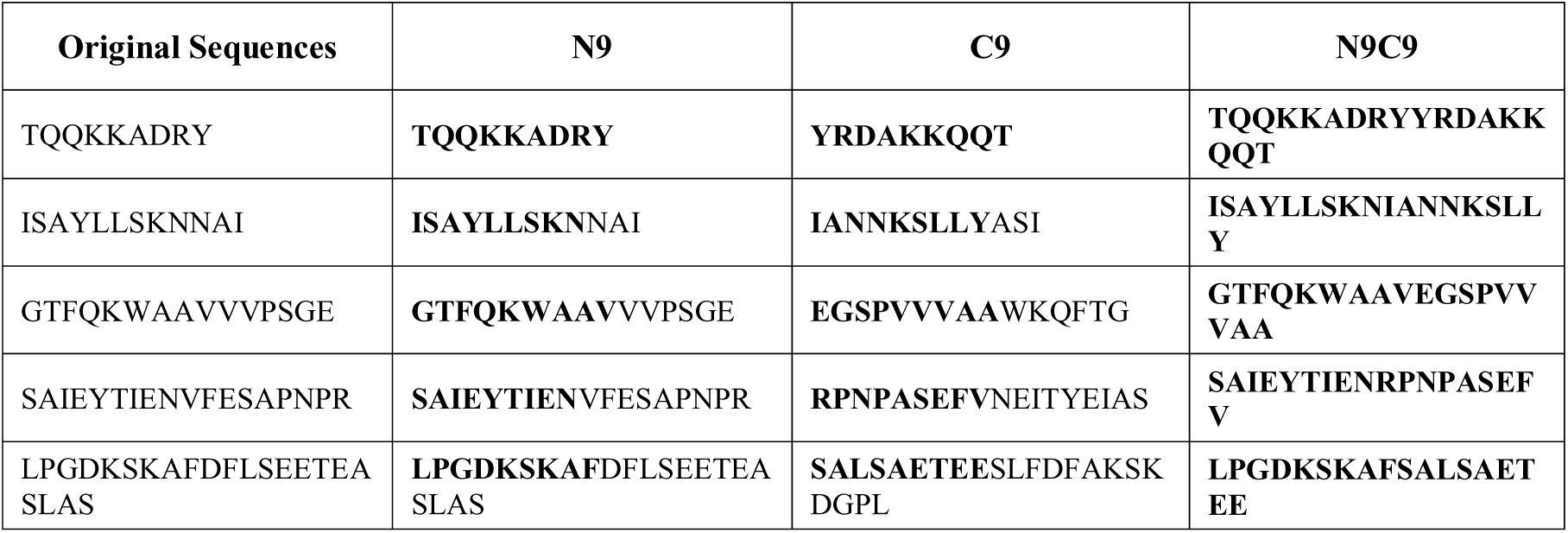
Generation of N_9_, C_9_, and N_9_C_9_ patterns from the original sequences with varying.

### length

Similarly, binary profile for pattern size with twenty-two residues (NC_22_) were also generated. The major challenge in calculating the binary profile for NC_22_ pattern was the varying length of the peptides. In order to tackle that situation, we have appended the dummy variable “X” in the sequences having length less than 22 as shown in Table 2.

**Table 2:**
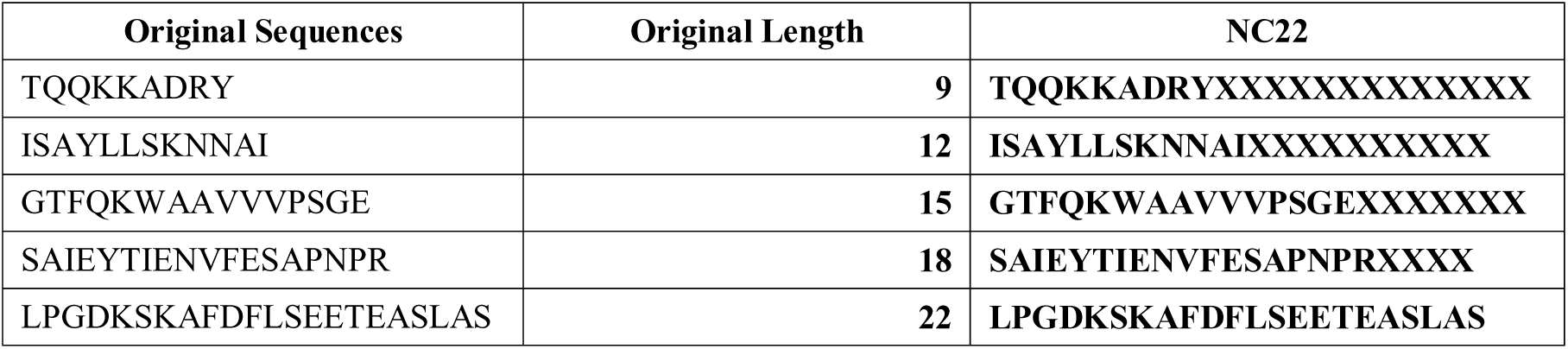
Generation of NC22 patterns from the original sequences with varying length.

### Model Development

In order to train and develop prediction models, we have utilized a diverse set of classifiers using scikit-learn [40] library of python such as decision tree (DT), random forest (RF), logistic regression (LR), extreme gradient boosting (XGB), k-nearest neighbor (KNN), gaussian naïve Bayes (GNB), extremely randomized tree (ET), and support vector classifier (SVC).

### Cross-Validation

To build more robust and accurate prediction models, we adopted the five-fold cross-validation technique that avoid overfitting and minimize bias in our generated models. Moreover, it also allows to assess the efficiency of the prediction models. As per the standard norms, we have implemented the five-fold cross validation technique on training dataset and kept the validation dataset untouched. As per the standard protocols, in this technique the entire dataset is divided into five parts, where four parts are used to train the model and tested on the remaining fifth one. The same process is iterated five times in such a way that each set/part gets the chance to act as testing dataset. Finally, the overall performance is the mean of the performances of five iterations.

### Evaluation of Parameters

To ensure the fair evaluation of the different generated models developed using various classifiers, we have used the well-established evaluation parameters. In this method, we implemented both threshold-dependent and threshold -independent parameters. For threshold-dependent parameters we consider sensitivity, specificity, accuracy, F1-score, kappa, and Mathews correlation coefficient (MCC). Meanwhile, area under the receiver operating characteristics curve (AUROC) is the measure of separability and it signifies how well the model is capable of distinguishing between the classes. Threshold dependent parameters were calculated using the following equations:

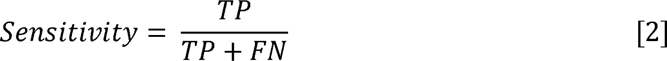

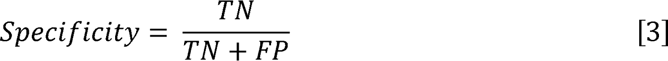

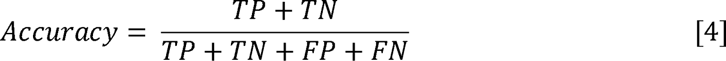

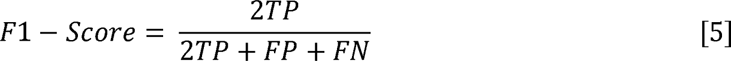

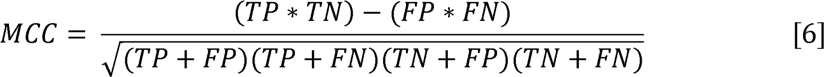

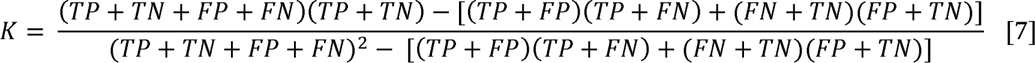

Where, TP stands true positive; TN stands for true negative; FP stands for false positive; FN stands for false negative

### Model Optimization

We evaluated eight different machine learning algorithms, and tuned the hyper parameters according to the training dataset. For this purpose, we used GridSearchCV to find the best performing model for each of our machine learning classifiers and optimised them by maximizing the AUROC.

### Similarity Search

In order to predict if the query peptide is a binder of a HLA-DRB1*04:01 using similarity search, we have implemented the Basic Local Alignment Search Tool (BLAST) [41] using the NCBI-blast executable version 2.13.0. We have created the custom database using our dataset by implementing “makeblastdb” module of NCBI-blast. Then, to make the prediction for query sequences we have implemented the “blastp” module with “blastp-short” as task since the peptide length are small. Top-hit against the query sequences were considered to assign the classes.

### Motif Analysis

To make the predictions using the small regions which are shared by all the sequences of a particular class also called motifs, we have implemented the Motif – Emerging and with Classes - Identification (MERCI) tool [42] with default the parameters. We have identified the motifs which are specific to the HLA-DRB1*04:01 binders and used them to assign the class as binder to the query/unseen data if the particular motif is found else assigned them as non-binders.

### Webserver Architecture

We have developed the user-friendly updated version of our old webserver HLA-DR4Pred, and named it as HLA-DR4Pred 2.0 to predict, scan, and design the HLA-DRB1*04:01 binding peptides. The front-end of the webserver was developed using HTML (v5), PHP (v7), CSS (v3), and JavaScript (v 1.8). The backend of the server uses Perl and python 3.6. The server compatibility is tested and confirmed to be compatible with all the modern devices including mobile, tablet, laptop, iMac, and desktop. The server is designed with six major modules including predict, scan, design, blast, motif-scan, and standalone to efficiently address the needs and tasks of various users.

## Results

### Composition Analysis

In the present study, we have calculated the average composition of each residue in HLA- DRB1*04:01 binders and non-binders in dataset. The amino acid composition is calculated using Pfeature [39]. The average residue composition for each dataset is provided in Figure 3, and it exhibits that serine residue is abundant in HLA-DRB1*04:01 binding peptides in comparison to the non-binding peptides. Moreover, the similarity in the trends of negative dataset generated randomly using Swiss-Prot [43] database and general proteome signifies that the negative dataset is not biased towards a particular amino acid or nature of amino acids.

**Figure 3:**
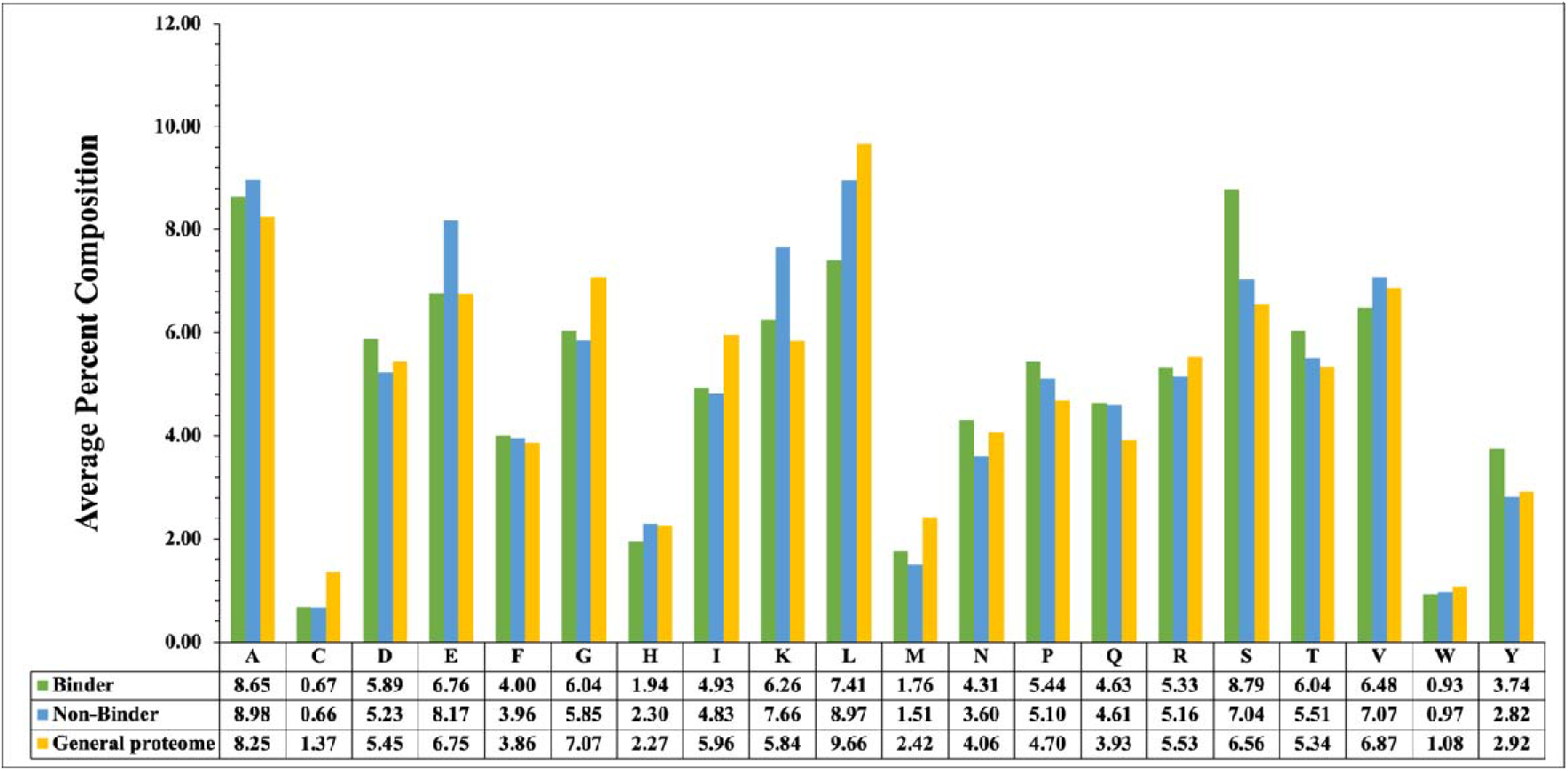
Average percent amino acid composition of HLA-DRB1*04:01 binder, non-binders and general proteome

### Position Preference Analysis

In this study, preference of particular residues at a specific positions in a peptide was studied by creating the “Weblogo” for HLA-DRB1*04:01 binders and non-binders in the main, and alternate dataset as shown in Figure 4. In case of HLA-DRB1*04:01 binders, positions 4, 5, and 6 are preferred by hydrophobic residues ‘L/F/Y/I/V’; where position 9 is covered by polar and uncharged amino acids ‘S/T/A’; position 10 is preferred by positive charged amino acid residues ‘K/R’; ‘S/A’ amino acids are favoured in positions 13-15; positions 16 and 17 are preferred by polar amino acids ‘S/T’, and ‘D’ residue is found to be most abundant at position 18 in HLA-DRB1*04:01 binding peptides. On the other hand, in case of HLA- DRB1*04:01 non-binding peptides, ‘P’ is preferred at position 2; positions 4, and 5, are most preferred by positive amino acid ‘K’; position 10 also preferred by positive charged residues ‘K/R’; and positions 14-19 showed abundance for residues ‘A/L’.

**Figure 4:**
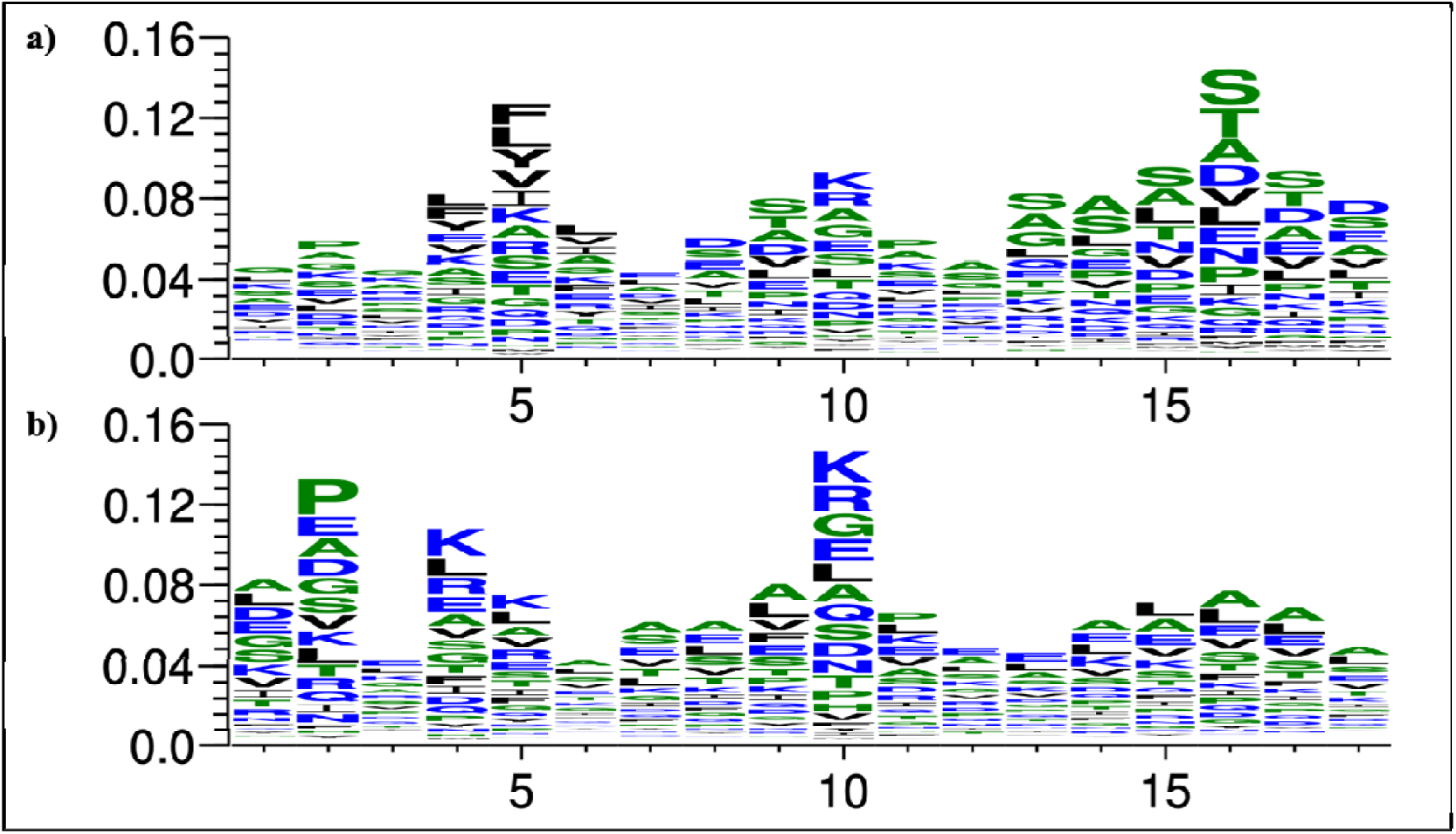
Positional preference representation using weblogo in a) HLA-DRB1*04:01 binders, b) HLA-DRB1*04:01 non-binders

### Performance of models on composition and binary profile based module

We have calculated the fifteen different types of composition features and the binary profiles for different patterns such as N_9_, C_9_, N_9_C_9_, and NC_22_ using the Pfeature module [39] to develop the prediction models using eight different classifiers from sklearn [40] library of python. The models were developed by implementing classifiers like DT, RF, LR, KNN, XGB, GNB, ET, and SVC. The models were trained on the training dataset and external validated on the validation dataset of main, and alternate dataset. The comprehensive performance of all the performing model developed on training and validation dataset using different types of features is reported in Supplementary Table 2. It was observed that ET classifier performs best among all other ML models. As shown in Figure 5, ET classifier based model developed on DPC features outperformed all the other models developed on other features, with AUROC of 0.90 on validation data of main dataset; and AUROC of 0.96 on validation data of alternate dataset. CTC based model performed second best with AUROC of 0.87 on validation data of main dataset. Although, binary profile pattern NC_22_ also outperformed the other patterns with AUROC of 0.87 on validation data of main dataset.

**Figure 5:**
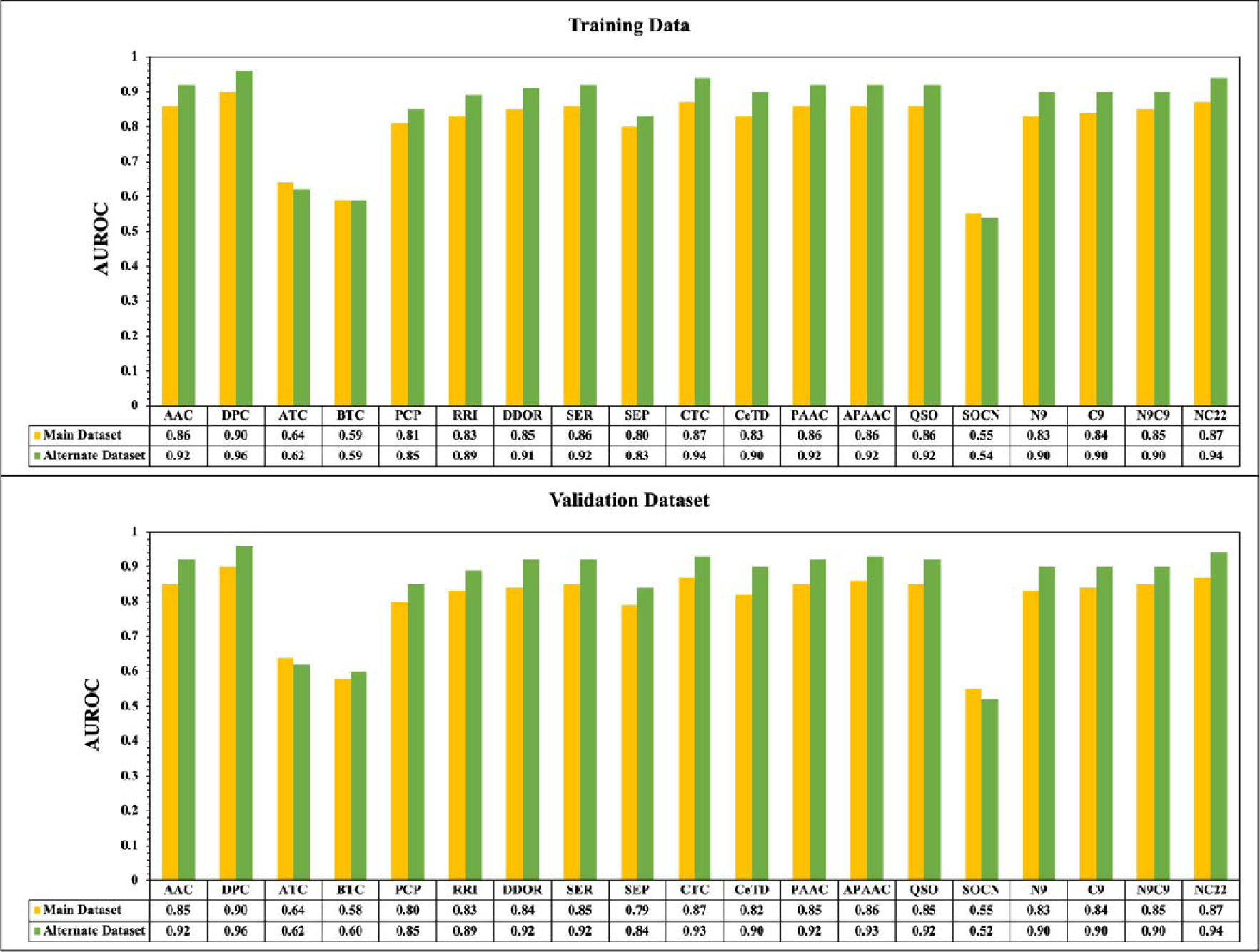
The performance of ET based classifier developed using fifteen different types of composition based features, and four different types binary profile based features on main, and alternate dataset.

### Performance of models on combined features

Further, we have combined all the features to develop the vector of size 2382 for each peptide belong to different datasets and develop the prediction models using eight different classifiers by hyper-tuning the parameters to maximize the AUROC on the training dataset and validation dataset. As shown in Figure 6, ET-based model developed using combined features outperformed all the other classifiers by attaining the maximum AUROC of 0.88 on validation data of main dataset. Supplementary Table 3 comprises the threshold-dependent and threshold-independent performance measure of all the classifiers on main, and alternate dataset.

**Figure 6:**
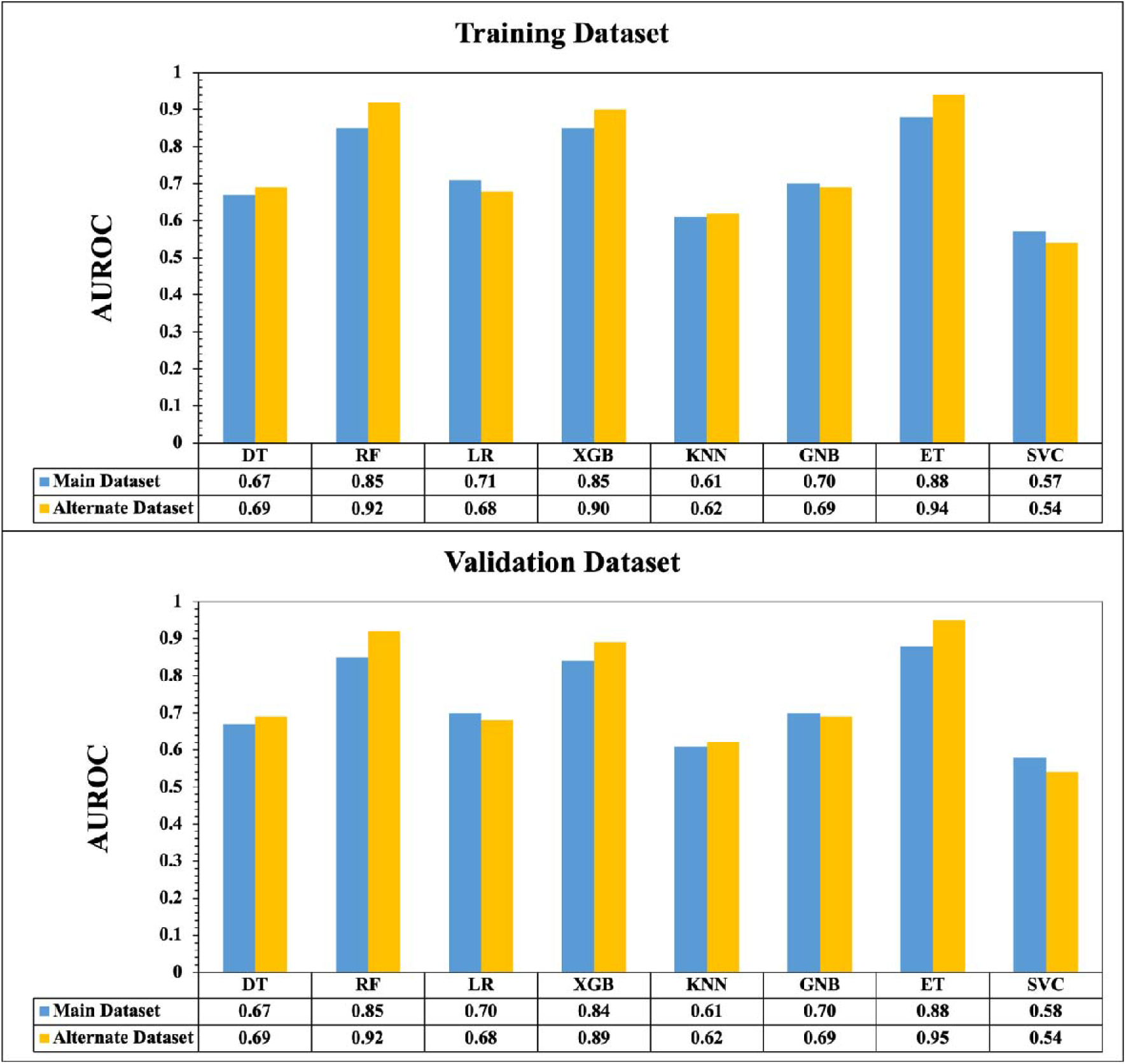
Performance for all model developed using various classifiers for, main, and alternate dataset on combined features on training and validation dataset

### Performance of models on selected features

In machine learning-based prediction methods, selecting an appropriate number of features to train the model is a critical step in order to reduce training time, avoid overfitting, and manage high-dimensional feature sets. In this approach, we have implemented the SVC-L1 (Support Vector Classifier with a linear kernel and L1 penalty regularization), which is highly efficient in identification of best-performing features in a sort span of time. Based on this technique, we left with 131 features for main dataset, and 258 features were selected in alternate dataset. Among all the classifiers, ET-based model has outperformed in each dataset with AUROC 0f 0.88 and 0.87 on training and validation of main dataset, and AUROC of 0.95 on both training and validation dataset of the realistic dataset as shown in Figure 7. Supplementary Table 4 contains the comprehensive results of all models developed using eight classifiers for each dataset.

**Figure 7:**
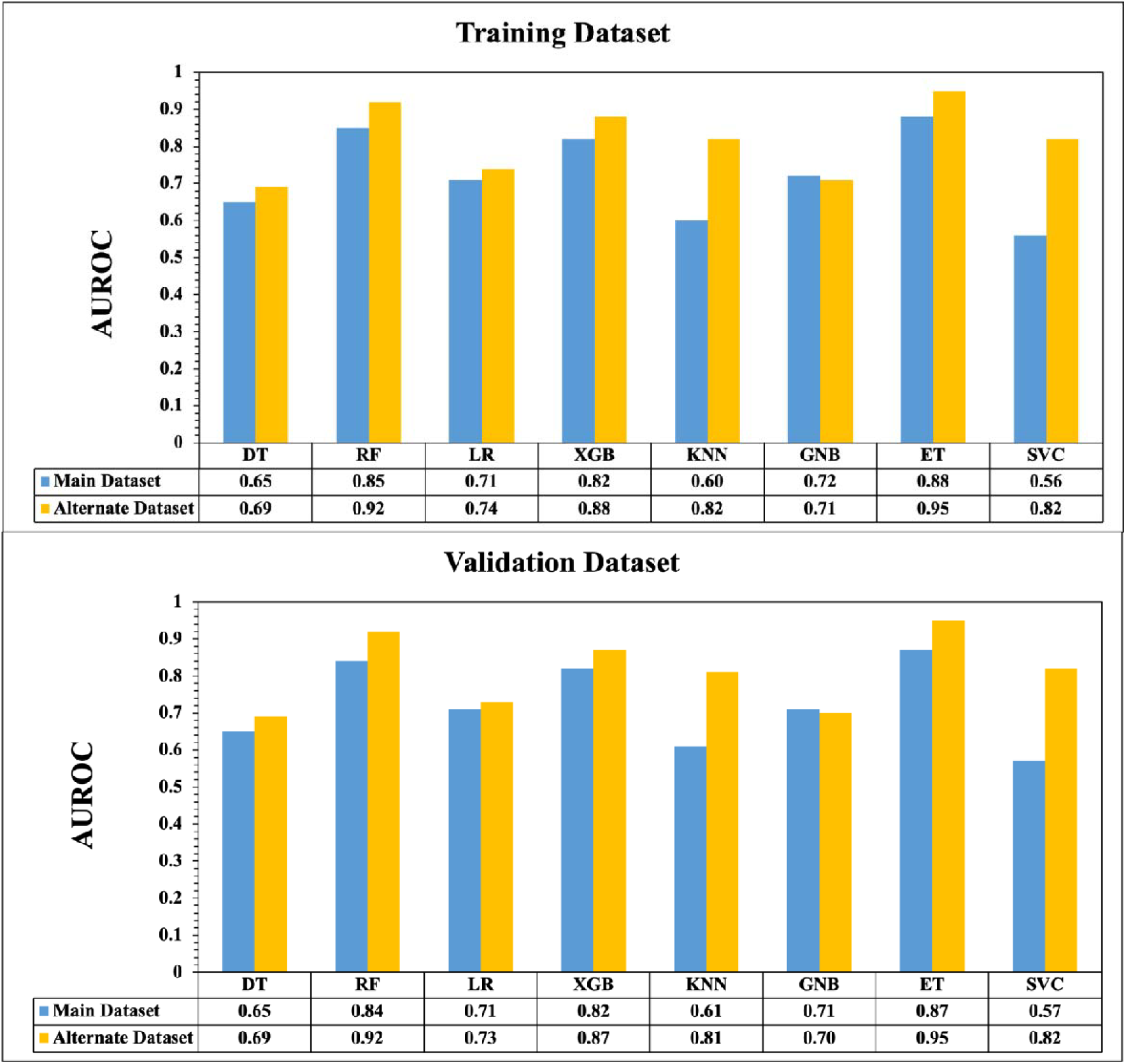
Performance for all model developed using various classifiers for main, and alternate dataset after reducing the features using SVC-L on training, and validation Dataset

### Performance of hybrid model

On observing the results of various machine learning classifiers on different type of features, it was found that ET-based model developed on DPC features outperformed all the other features with AUROC of 0.90 on validation data of main dataset. Consequently, in order to enhance the performance, we have integrated the ET-based model of DPC with similarity search using BLAST [41], and call it the hybrid model. We have implemented BLAST with varying e-value in order to find the optimal value at which we can achieve the maximum AUROC. Table 3 captures the results for each dataset at different e-values for training as well as validation dataset. We varied the e-value from e^-6^ to e^2^ and main dataset is able to achieve AUROC 0.93 on validation dataset at e-value 1.0, followed by alternate dataset achieves the AUROC of 0.99 on validation dataset. Comprehensive results are provided in Supplementary Table 5. The same model has been implemented at the backend of the server and respective standalone packages.

**Table 3:** Performance of hybrid model at different e-values for main, and alternate dataset on training and validation dataset.

| E-value | Dataset | Main Dataset |  |  | Alternate Dataset |  |  |
| --- | --- | --- | --- | --- | --- | --- | --- |
|  |  | Acc | AUROC | MCC | Acc | AUROC | MCC |
| **1.00E-06** | **Training** | 82.19 | 0.9 | 0.64 | 88.99 | 0.95 | 0.64 |
|  | **Validation** | 81.79 | 0.9 | 0.64 | 89.58 | 0.95 | 0.65 |
| **1.00E-05** | **Training** | 82.25 | 0.9 | 0.65 | 88.35 | 0.95 | 0.63 |
|  | **Validation** | 82.01 | 0.91 | 0.64 | 89.36 | 0.95 | 0.65 |
| **1.00E-04** | **Training** | 82.38 | 0.9 | 0.65 | 88.86 | 0.95 | 0.63 |
|  | **Validation** | 82.36 | 0.91 | 0.65 | 90.14 | 0.95 | 0.66 |
| **1.00E-03** | **Training** | 82.68 | 0.91 | 0.65 | 88.64 | 0.95 | 0.63 |
|  | **Validation** | 82.67 | 0.91 | 0.65 | 90.13 | 0.96 | 0.67 |
| **1.00E-02** | **Training** | 83.22 | 0.91 | 0.66 | 88.57 | 0.96 | 0.63 |
|  | **Validation** | 83.21 | 0.92 | 0.67 | 90.17 | 0.96 | 0.67 |
| **1.00E-01** | **Training** | 84.57 | 0.92 | 0.69 | 88.4 | 0.96 | 0.63 |
|  | **Validation** | 85.02 | 0.93 | 0.7 | 90.14 | 0.96 | 0.67 |
| **1.00E+00** | **Training** | **84.77** | **0.92** | **0.7** | **88.4** | **0.98** | **0.63** |
|  | **Validation** | **85.31** | **0.93** | **0.71** | **90.14** | **0.99** | **0.67** |
| **1.00E+01** | **Training** | 84.82 | 0.92 | 0.7 | 88.44 | 0.96 | 0.63 |
|  | **Validation** | 86.14 | 0.93 | 0.72 | 89.42 | 0.96 | 0.65 |
| **1.00E+02** | **Training** | 84.26 | 0.92 | 0.69 | 89.59 | 0.96 | 0.66 |
|  | **Validation** | 85 | 0.93 | 0.7 | 89.83 | 0.96 | 0.66 |

\*Sens: Sensitivity; Spec: Specificity; Acc: Accuracy; AUROC: Area under the receiver operating characteristic curve; F1: F1 score; MCC: Matthews Correlation Coefficient; K: Cohen's Kappa

### Motif analysis

In this study, we have implemented MERCI software with default parameters to obtain the specific regions i.e. motifs from the main dataset which are highly specific to HLA- DRB1*04:01 binders but absent in non-binder, similar procedure was repeated for non-binders where we searched for non-binder specific motifs which are exclusively present in the binders and absent in the binder sequences. In Table 4, we have reported motifs specific to binders and non-binders along with their coverage in the positive and negative dataset. Residue T, V, F, P, Q, and T are dominant in binders, where residue D, Y, and K covers the most of the motifs.

**Table 4:** Exclusive motifs specific to HLA-DRB1*04:01 binder and non-binders.

| HLA-DRB1*04:01 Binders |  |  | HLA-DRB1*04:01 Non-binders |  |  |
| --- | --- | --- | --- | --- | --- |
| Motif | # Sequences | Coverage | Motif | # Sequences | Coverage |
| A-F-V-K-D | 56 | 56 | Y-D-G-K-D | 335 | 335 |
| V-A-F-V-K-D | 53 | 109 | Y-D-G-K-D-Y | 315 | 650 |
| D-V-A-F-V-K-D | 49 | 158 | A-Y-D-G-K-D | 293 | 943 |
| F-T-P-E-T | 46 | 204 | R-K-W-E-A | 276 | 1219 |
| F-T-P-E-T-N | 46 | 250 | A-Y-D-G-K-D-Y | 274 | 1493 |
| F-T-P-E-T-N-P | 46 | 296 | R-K-W-E-A-A | 251 | 1774 |
| T-P-E-T-N | 46 | 342 | S-D-H-E | 249 | 1993 |
| T-P-E-T-N-P | 46 | 388 | Y-D-G-K-D-Y-I | 243 | 2236 |
| D-Q-T-V-I | 45 | 433 | Y-D-G-K-D-Y-I-A | 236 | 2472 |
| D-Q-T-V-I-Q | 45 | 478 | Y-D-G-K-D-Y-I-A-L | 236 | 2708 |
| F-V-K-D-Q | 45 | 523 |  |  |  |
| F-V-K-D-Q-T | 45 | 568 |  |  |  |
| F-V-K-D-Q-T-V | 45 | 613 |  |  |  |
| F-V-K-D-Q-T-V-I | 45 | 658 |  |  |  |
| F-V-K-D-Q-T-V-I-Q | 45 | 703 |  |  |  |
| I-F-T-P-E | 45 | 748 |  |  |  |
| I-F-T-P-E-T | 45 | 793 |  |  |  |
| I-F-T-P-E-T-N | 45 | 838 |  |  |  |
| I-F-T-P-E-T-N-P | 45 | 883 |  |  |  |
| K-D-Q-T-V-I | 45 | 928 |  |  |  |
| K-D-Q-T-V-I-Q | 45 | 973 |  |  |  |
| S-I-F-T-P | 45 | 1018 |  |  |  |
| S-I-F-T-P-E | 45 | 1063 |  |  |  |
| S-I-F-T-P-E-T | 45 | 1108 |  |  |  |
| S-I-F-T-P-E-T-N | 45 | 1153 |  |  |  |
| S-I-F-T-P-E-T-N-P | 45 | 1198 |  |  |  |
| V-K-D-Q-T | 45 | 1243 |  |  |  |
| V-K-D-Q-T-V | 45 | 1288 |  |  |  |
| V-K-D-Q-T-V-I | 45 | 1333 |  |  |  |
| V-K-D-Q-T-V-I-Q | 45 | 1378 |  |  |  |

### Comparison with the available methods

It is important to evaluate the newly developed method’s effectiveness in comparison to that of existing methods in order to gain insights into the pros and cons of a newly developed method. Since, HLADR4Pred2.0 is an update of HLADR4Pred [32], hence its comprehensive comparison is required to understand the advantages of the newer version over older versions. Supplementary Table 6 accumulates differences in the HLADR4Pred and HLADR4Pred2.0 at the level of dataset, implemented features, prediction approach, webserver and standalone. Other than HLADR4Pred, there are number of other methods with the ability to predict the binders for HLA-class II alleles. Hence, it is crucial to benchmark the performance of the other existing methods with HLADR4Pred2.0. For that purpose, we have taken out the validation dataset and tested the performance of the existing methods on the same. Propred [33] is able to predict the HLA-DR binding sites and able to achieve 55.26% accuracy with AUROC 0.74, where NetMHCIIpan 4.0 [35,36] achieved accuracy 65.82% with AUROC 0.72, followed by TEPITOPE [44] with accuracy of 67.75% with balanced sensitivity and specificity, SMM-align [45] predicts the MHC class II binding affinity using stabilization matrix alignment method achieved accuracy of 67.95%. Artificial neural network based method i.e. NNAlign [34] develop the model on sequence motifs detected in the training data, attained the accuracy of 68.64%, followed by consensus IEDB method with uses the consensus of SMM-Align, NNAlign, and Sturniolo method to calculate the adjusted rank based on which the predictions are made and it attained the accuracy of 69.41% on the independent dataset. Finally, older version of HLADR4Pred2.0 achieved the accuracy of 75.04 with AUROC 0.69, but the difference between sensitivity and specificity is significant. Our new approach has outperformed all the existing with methods with AUROC of 0.93 and accuracy 85.31%. These results in Table 5 showed that HLADR4Pred2.0 is an reliable method which has outperformed the other methods on the independent validation dataset of main dataset which was not used while training or testing the model.

**Table 5:** Comparison of HLADR4Pred2.0 approach with the existing methods on validation dataset of main dataset.

| Methods | Sensitivity | Specificity | Accuracy | AUROC | F1-score | Kappa | MCC |
| --- | --- | --- | --- | --- | --- | --- | --- |
| **Propred** | 78.378 | 44.156 | 55.263 | 0.735 | 0.532 | 0.181 | 0.219 |
| **NetMHCIIpan 4.0** | 65.249 | 66.253 | 65.819 | 0.717 | 0.623 | 0.311 | 0.313 |
| **TEPITOPE** | 68.278 | 67.336 | 67.747 | NA | 0.297 | -0.347 | 0.353 |
| **SMM-Align** | 68.535 | 67.495 | 67.948 | NA | 0.292 | -0.357 | 0.358 |
| **NNAlign** | 68.946 | 68.410 | 68.643 | NA | 0.288 | -0.367 | 0.371 |
| **Consensus IEDB** | 69.409 | 69.404 | 69.406 | NA | 0.283 | -0.380 | 0.385 |
| **Hladr4pred** | 54.098 | 79.447 | 75.036 | 0.690 | 0.430 | 0.279 | 0.289 |
| **HLADR4Pred2.0** | 88.32 | 82.30 | 85.31 | 0.93 | 0.86 | 0.71 | 0.71 |

\*Sens: Sensitivity; Spec: Specificity; Acc: Accuracy; AUROC: Area under the receiver operating characteristic curve; F1: F1 score; MCC: Matthews Correlation Coefficient; K: Cohen's Kappa

### Case Study: HLA-DRB1*04:01-binders in COVID-19 variants

Recent studies report that HLA-DRB1*04:01 binding sites are associated with the severity of COVID-19 patients [46–48]. The mutations associated with spike protein in COVID-19 variants can alter the binding of peptides [49,50]. In order to understand the effect of mutations in different variants of COVID-19 with the HLA-binding peptides, we utilized “SCAN” module of our HLA-DR4Pred 2.0 server (https://webs.iiitd.edu.in/raghava/hladr4pred2/scan.php). First we created mutated proteins of COVID-19 variants using the reference spike protein sequence. As reported in Centres for Disease Control and Prevention (CDC portal) (https://www.cdc.gov/hai/data/portal/), the alpha variant possess seven mutation named as N501Y, A570D, D613G, P681H, T716I, D981A and D1118H, whereas beta variant in corporate D80A, D215G, K417N, E484K, N501Y, D614G, A701V, L18F and R246I mutations. Similarly, spike protein of delta variant incorporates T19R, T95I, G142D, R158G, L452R, T478K, D614G, L681R and D950N mutations. Recently, reported COVID-19 variant Omicron possess highest number of mutations i.e., 30 mutations in spike protein A67V, del 69-70, T95I, G142D, del 143-145, del 211, L212I, ins214EPE, G339D, S371L, S373P, S375F, K417N, N440K, G446S, S477N, T478K, E484A, Q493K, G496S, Q498R, N501Y, Y505H, T547K, D614G, H655Y, N679K, P681H, N764K, D796Y, N856K, Q954H, N969K, L981F. Currently, we created the mutated proteins of different variants of COVID-19 and predict the binding peptides and effect of mutation on bindings in different COVID variants. We observed that in alpha variant (D981A and D613G), beta variant (D80A), gamma variant (D137Y), delta variant (G142D, L681R) and omicron associated mutations alter the nature of HLA-binding peptides to non-binders or vice versa, as shown in Table 6.

**Table 6:**
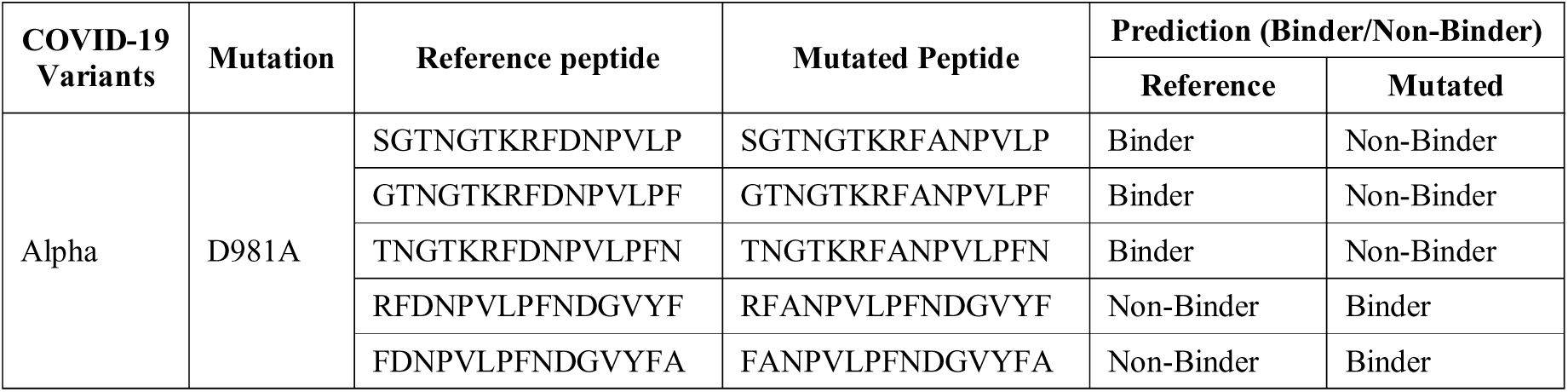
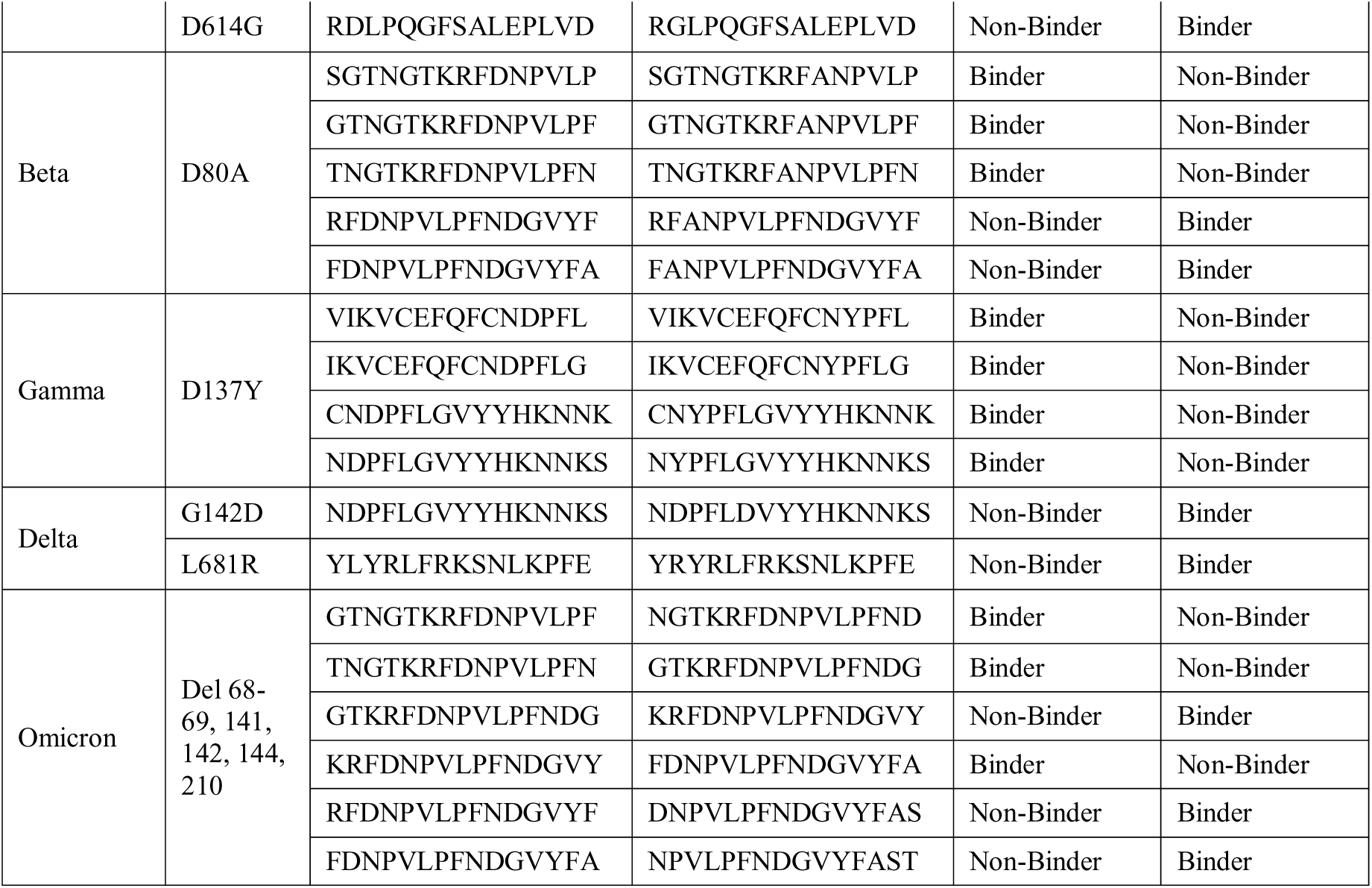
Alterations in the binding peptides of HLA-DRB1*04:01 by mutations in Spike protein of SARS-CoV-2 variants.

### Webserver Implementation

We created the user-friendly webserver "HLADR4Pred 2.0" to assist the scientific community available at URL https://webs.iiitd.edu.in/raghava/hladr4pred2. There are six major modules in the server such as, “PREDICT”, “SCAN”, “DESIGN”, “BLAST”, “MOTIF-SCAN”, and “STANDALONE”. The description of each module in provided below. The module “PREDICT” allows users to predict the potential of an uncharacterized peptide to a HLA-DRB1*04:01 binder. The module “SCAN” allows user to provide sequences with length more than 22, which is a constraint in the predict module. In this module, users are asked to choose a desired window size on which the overlapping patterns are generated from the input sequence(s) and used them to make predictions. The module “DESIGN” permits users to generate all the possible mutants of an input sequence by mutating each residue at a time and use the same to predict if the mutated pattern is a binder. In the “BLAST” module, user can make the predictions if a submitted sequence(s) is a binder or non-binder by performing similarity search using BLAST. In the module “MOTIF-SCAN” HLA-DRB1*04:01 binding motifs are searched in the input sequences and the predicted as binder if the motifs is found else designated as non-binder.

### Discussion and Conclusion

HLA system is the major histocompatibility complex in humans and is the most import part of our immune system [6]. HLA genes regulate the immune responses while infectious diseases and viral/pathogenic attack and provide protections [1,51–53]. Due to high polymorphism, thousands of HLA alleles are reported in IMGT/HLA database [5], out of which few were associated with number of diseases. In the past few decades researches proves that one of the HLA-DR4 family allele HLA-DRB1*04 play major role in the regulation of immune responses and associated with several autoimmune disorders and COVID-19 severity [46,47,51,54]. In the current study, we have extracted experimentally validated 12676 positive dataset (i.e. HLA-DRB1*04:01 binding peptides and 86300 negative dataset (non-binding peptides) from IEDB. Firstly, we compute composition based, and binary profile based features using Pfeature modules. We have developed various machine learning models using eight different classifiers such as SVC, DT, RF, XGB, KNN, LR, ET, and GNB. As shown in most of results developed on different feature in datasets, ET based models outperform the other classifiers. After selecting the best features we obtained that DPC based models achieves the highest AUROC of 0.90 on main dataset, followed by AUROC of 0.96 on alternate dataset with accuracy is more than 82% on training and validation datasets (Supplementary Table 2). In order to achieve the maximum performance we have merged machine learning technique with similarity search using BLAST and attained AUROC 0.93 on validation dataset of main dataset. To compare our performance of our model with the existing methods, we have considered seven different methods as shown in Table 5, and found out that HLA-DR4Pred2.0 has outperformed all the other methods with AUROC of 0.93. In order to serve the scientific community we have developed a webserver and standalone package using the best features and classifiers. HLA- DR4Pred 2.0 incorporates five modules such as PREDICT, SCAN, DESIGN, BLAST, and MOTIF-SCAN. HLA-DR4Pred 2.0 tool predict the binding or non-binding peptides for MHC-Class II allele HLA-DRB1*04:01. Our webserver is freely accessible at https://webs.iiitd.edu.in/raghava/hladr4pred2/ and standalone package is available at https://webs.iiitd.edu.in/raghava/hladr4pred2/standalone.php.

### Funding Source

The current work has been supported by the Department of Biotechnology (DBT) grant BT/PR40158/BTIS/137/24/2021.

### Conflict of interest

The authors declare no competing financial and non-financial interests.

### Authors’ contributions

SP, AD, and GPSR collected and processed the datasets. SP, AD, and GPSR implemented the algorithms and developed the prediction models. AD, SP, and GPSR analysed the results. SP created the back-end and front-end user interface of the web server. SP, NK, AD, and GPSR performed the writing, reviewing and draft preparation of the manuscript. GPSR conceived and coordinated the project. All authors have read and approved the final manuscript.

## Acknowledgements

Authors are thankful to the University Grants Commission (UGC), and Department of Biotechnology (DBT) for fellowships and financial support, and the Department of Computational Biology, IIITD New Delhi for infrastructure and facilities. We would like to acknowledge that Figures were created using BioRender.com.

## Data Availability Statement

All the datasets used in this study are available at the “HLA-DR4Pred 2.0” web server, https://webs.iiitd.edu.in/raghava/hladr4pred2/dataset.php.

